# Gene transfers, like fossils, can date the Tree of Life

**DOI:** 10.1101/193813

**Authors:** Adrián A. Davín, Eric Tannier, Tom A. Williams, Bastien Boussau, Vincent Daubin, Gergely J. Szöllősi

## Abstract

Biodiversity has always been predominantly microbial and the scarcity of fossils from bacteria, archaea and microbial eukaryotes has prevented a comprehensive dating of the tree of life. Here we show that patterns of lateral gene transfer deduced from the analysis of modern genomes encode a novel and abundant source of information about the temporal coexistence of lineages throughout the history of life. We use new phylogenetic methods to reconstruct the history of thousands of gene families and demonstrate that dates implied by gene transfers are consistent with estimates from relaxed molecular clocks in Bacteria, Archaea and Eukaryotes. An inspection of discrepancies between transfers and clocks and a comparison with mammal fossils show that gene transfer in microbes is potentially as informative for dating the tree of life as the geological record in macroorganisms.

---

Until Zuckerkandl and Pauling put forth the “molecular clock”^1^ hypothesis, the geological record alone provided the timescale for evolutionary history. Their demonstration that distances between amino acid sequences correlate with divergence times estimated from fossils demonstrated that information in DNA can be used to date the Tree of Life. Since then, the theory and methodology of the molecular clock have been developed extensively, and inferences from clock analyses (such as the diversification of placentals before the demise of dinosaurs^2,3^) hotly debated. Despite these controversies, combining information from rocks and clocks is now widely accepted to be indispensable^3,4,5^: state-of-the-art estimates of divergence times rely on sequence based relaxed molecular clocks anchored by multiple fossil calibrations. This approach provides information on both the absolute timescale and the relative variation of the evolutionary rates across the phylogeny (Fig.1a). Yet, because most life is microbial, and most microbes do not fossilize, major uncertainties remain about the ages of microbial groups and the timing of some of the earliest and most important events in life’s evolutionary history^6,7^.

In addition to leaving only a faint trail in the geological record, the evolution of microbial life has also left a tangled phylogenetic signal due to extensive lateral gene transfer (LGT). LGT, the acquisition of genetic material potentially from distant relatives, has long been considered an obstacle for reconstructing the history of life^8^, because different genetic markers can yield conflicting estimates of the species phylogeny. However, it has been previously shown that transfers identified using appropriate phylogenetic methods carry information that can be harnessed to reconstruct species history^9–14^. This is possible because different hypotheses of species relationships yield different LGT scenarios and can thus be evaluated using phylogenetic models of genome evolution^15–19^. But in addition to carrying information about the relationships among species, transfers should also carry a record of the timing of species diversification because they have occurred between species that existed at the same time^20–22^. As a consequence, a transfer event can be used to establish a relative age constraint between nodes in a phylogeny independently of any molecular clock hypothesis: the ancestor node of the donor lineage must predate the descendant node of the receiving lineage (Fig.1b, Fig.S8). Below we show that the dating information carried by transfers is consistent with molecular clock based estimates of relative divergence times in representative groups from the three domains of life.

**Figure 1.**
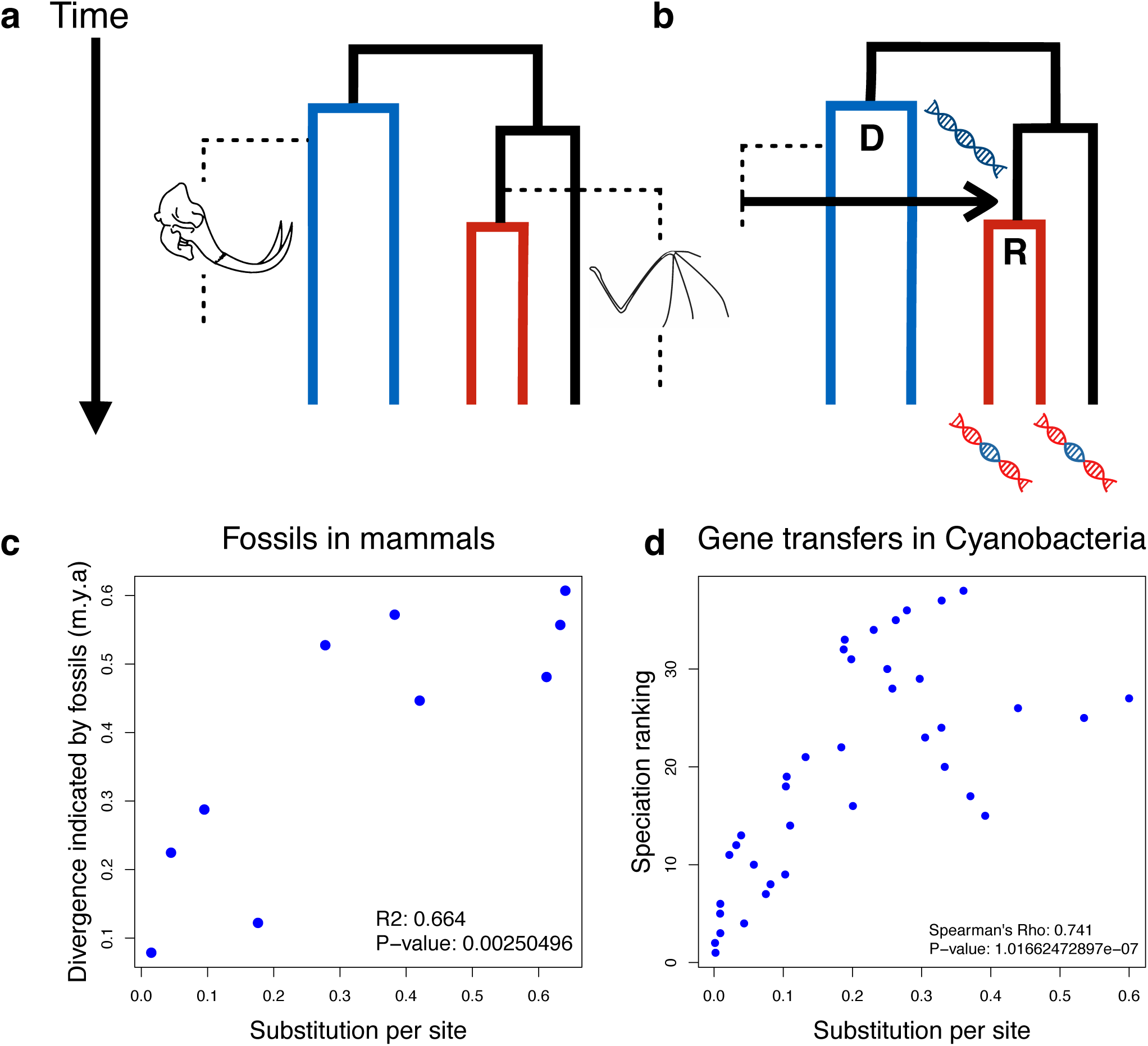
Gene transfers, like fossils, carry information on the timing of species divergence: **a)** The geological record provides the only source of information concerning absolute time: the age of the oldest fossil representative of a clade provides direct evidence on its minimum age, but inferring maximum age constraints (e.g. dashed line for the red clade), and by extension the relative age of speciation nodes, must rely on indirect evidence on the absence of fossils in the geological record^5,23–25^. **b)** Gene transfers, in contrast, do not carry information on absolute time, but they do define relative node age constraints by providing direct evidence for the relative age of speciation events: the gene transfer depicted by the black arrow implies that the diversification of the blue donor clade predates the diversification of the red clade (*i.e.* node D is necessarily older than node R). **c)** Sequence divergence (here measured in units of expected number of nucleotide substitutions along a strict molecular clock time tree, see supplementary materials) for 36 mammals^2^ is correlated (Pearson’s R^2^=0.664, p<10^-2^) with age estimates based on the fossil record (ages corresponding to the time of divergence in million years). **d)** A similar relationship can be seen for gene transfer based relative ages by plotting the sequence divergence (measured similar to part c) against the relative age of ancestral nodes for 40 cyanobacterial genomes (Spearman’s rank correlation rho=0.741, p<10^-6^) inferred by the MaxTiC (**max**imal **ti**me **c**onsistency) algorithm^26^.

We examined genome-scale datasets consisting of homologous gene families from complete genomes in Cyanobacteria (40 genomes^27^), Archaea (60 genomes^28^) and Fungi (60 genomes^29^). For each gene family we used the species tree-aware probabilistic gene tree inference method ALE undated^27,30^ to sample evolutionary scenarios involving events of duplication, transfer and loss of genes conditional on a rooted species phylogeny and the multiple sequence alignment of the family. The undated reconciliation method does not impose any constraint on possible donor-recipient branch pairs aside from forbidding transfers to go from descendants to parents (Fig. S9). For putative gene transfer events, we recorded the donor and recipient branches and used the frequency with which they occurred among the sampled scenarios to filter transfers and weigh the relative age information they imply. Because the reference species tree is not dated, individual transfers can imply conflicting information about the relative age of speciation nodes (Fig. S11). To extract a maximal subset of transfers consistent with each other, we used the newly developed optimization method MaxTiC^26^ (**m**aximal **t**ime **c**onsistency, see also supplementary text). A maximal subset of consistent transfers specifies a time order of speciation events in the species tree. For instance, using MaxTiC on the 4816 transfers that correspond to relative age constraints (see Figs 1b, S8, S10) in the 5322 gene families considered for Cyanobacteria, we identified a maximal subset of 3322 (69%) transfers that are consistent (Table S1). This maximal subset of transfers implies a time order of speciations that correlates with the distance between amino acid sequences of extant organisms (Spearman’s ρ = 0.741; p < 10^-6^; Fig. 1d, S9). A similar correlation (Fig. 1c) can be observed if, following Zuckerkandl and Pauling^1^, we compare fossil dates and sequence divergence in mammals^2^ (10 time points, Pearson’s R^2^=0.664; p = 0.0025 and Spearman’s ρ = 0.83; p = 0.0056).

We observed a strong correlation between time estimates from MaxTiC and molecular clocks in all our datasets (p<10^-3^ - Fig S14-S16). This suggests that LGT indeed carries information on the relative age of nodes in all three domains of life. However, it is not conclusive because part of the correlation trivially results from the fact that parent nodes are necessarily both older and more distant to extant sequences than their direct descendants^31^. To control for this effect, we compared the relative time orders of speciation events inferred from transfers to dates obtained using molecular clocks in the absence of calibrations. We used Phylobayes^32^ on a concatenate of nearly universal gene family alignments to sample chronograms (*i.e.*, dated trees) under four different uncalibrated molecular clock models^33^ (the strict molecular clock, the autocorrelated lognormal, the uncorrelated gamma and the white-noise model). As a control for the shape of the tree, we measured the random expectation by sampling chronograms from the prior on divergence times but keeping the species phylogeny fixed (without any sequence information). To compare the dating information from transfers to the information conveyed by fossils, we used the same uncalibrated approach on the same mammalian dataset as above^2,34^ and derived relative node age constraints from fossil calibrations (see supplementary text). For the prokaryotic and fungi datasets, we derived relative node age constraints from the maximal consistent subsets of transfers obtained using MaxTiC^26^. For both fossil- and transfer-based constraints, we then measured the fraction of constraints that are in agreement with each chronogram. As Fig. 2 shows, both fossil- and transfer-based constraints agree with uncalibrated molecular clocks significantly more than expected by chance. The observed agreement is robust to the choice of different clock models (Fig. 2), priors on divergence time and models of protein evolution (Figs. S17-S19). This result demonstrates the presence of genuine and substantial dating signal in gene transfers.

**Figure 2.**
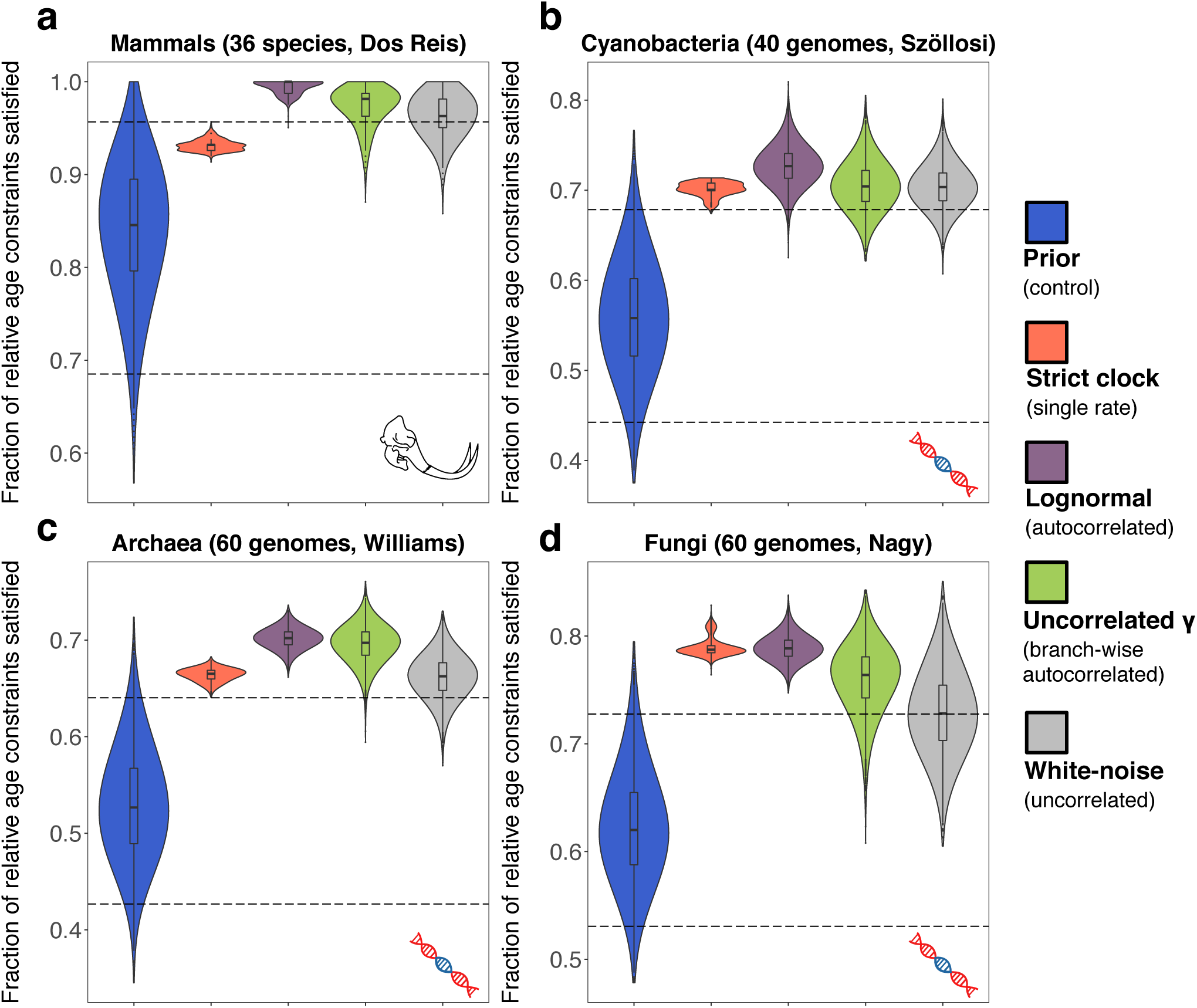
Agreement between transfer based relative ages and molecular clocks: **a)** Relative ages derived from 12 fossil calibrations from a phylogeny of 36 extant mammals were compared with node ages sampled from four different relaxed molecular clock models implemented in Phylobayes and with node ages derived from random chronograms, keeping the species phylogeny fixed. **b-d)** Relative ages derived from gene transfers using the MaxTiC algorithm were compared with estimates from the same 5 models as in a). For each model and each sampled chronogram we calculated the fraction of relative age constraints that are satisfied. On each plot, we show the distribution of the fraction of relative age constraints satisfied by 5000 sampled chronograms. The blue distribution corresponds to chronograms drawn from the prior with the 95% confidence interval denoted by dashed lines, orange to the strict molecular clock, purple to the autocorrelated lognormal, green to the uncorrelated gamma and grey to the white-noise models.

Interestingly, molecular clock models show differences in their agreement with relative time constraints. As expected, the strict molecular clock model generally explores a narrow range of dated trees compared to relaxed clocks. However, on average, chronograms based on the strict molecular clock agree less with relative time constraints than those based on relaxed clock models. This is particularly clear in mammals, where the median fraction of satisfied constraints falls within the 95% confidence interval of the random control (Fig. 2a). This is caused, in large part, by the accelerated evolutionary rate in rodents being interpreted (in the absence of fossil calibrations) as evidence for an age older than that implied by fossils (Fig. S4). The lognormal model is best suited to recover such autocorrelated (*e.g.* clade specific) rate variations along the tree, and indeed exhibits a median of 100% agreement with fossil based relative age constraints. The uncorrelated gamma model performs second best, perhaps because it is, in fact, autocorrelated along each branch^34^. Consistent with this idea, the completely uncorrelated white noise model fares worst (Fig. 2a-d). This is in agreement with previous model comparisons in eukaryotes, vertebrates and mammals^34^. A similar pattern is apparent when considering LGT-derived relative age constraints in Cyanobacteria, Archaea and Fungi, suggesting strong autocorrelated variation of evolutionary rates in these groups that are best recovered by the lognormal model (Fig. 2b-d).

The motivating principle of the MaxTiC algorithm is that transfers from the maximum consistent set carry a robust and genuine dating signal, while conflicting transfers are likely artefactual. Two lines of evidence suggest that this is indeed the case: first, the agreement of relative time constraints derived from transfers excluded by MaxTiC with the node ranking inferred by uncalibrated molecular clocks tends to be lower than random (Fig. S12). Second, while the average sequence divergences for donor clades tend to be higher than for corresponding recipient clades in the set of self-consistent transfers (p < 10^-8^ one sided T-test for difference greater than zero, see Fig. 3.), they are lower for those discarded by MaxTiC (p < 10^-8^ one sided T-test for difference lower than zero, cf. Fig. 3).

**Figure 3.**
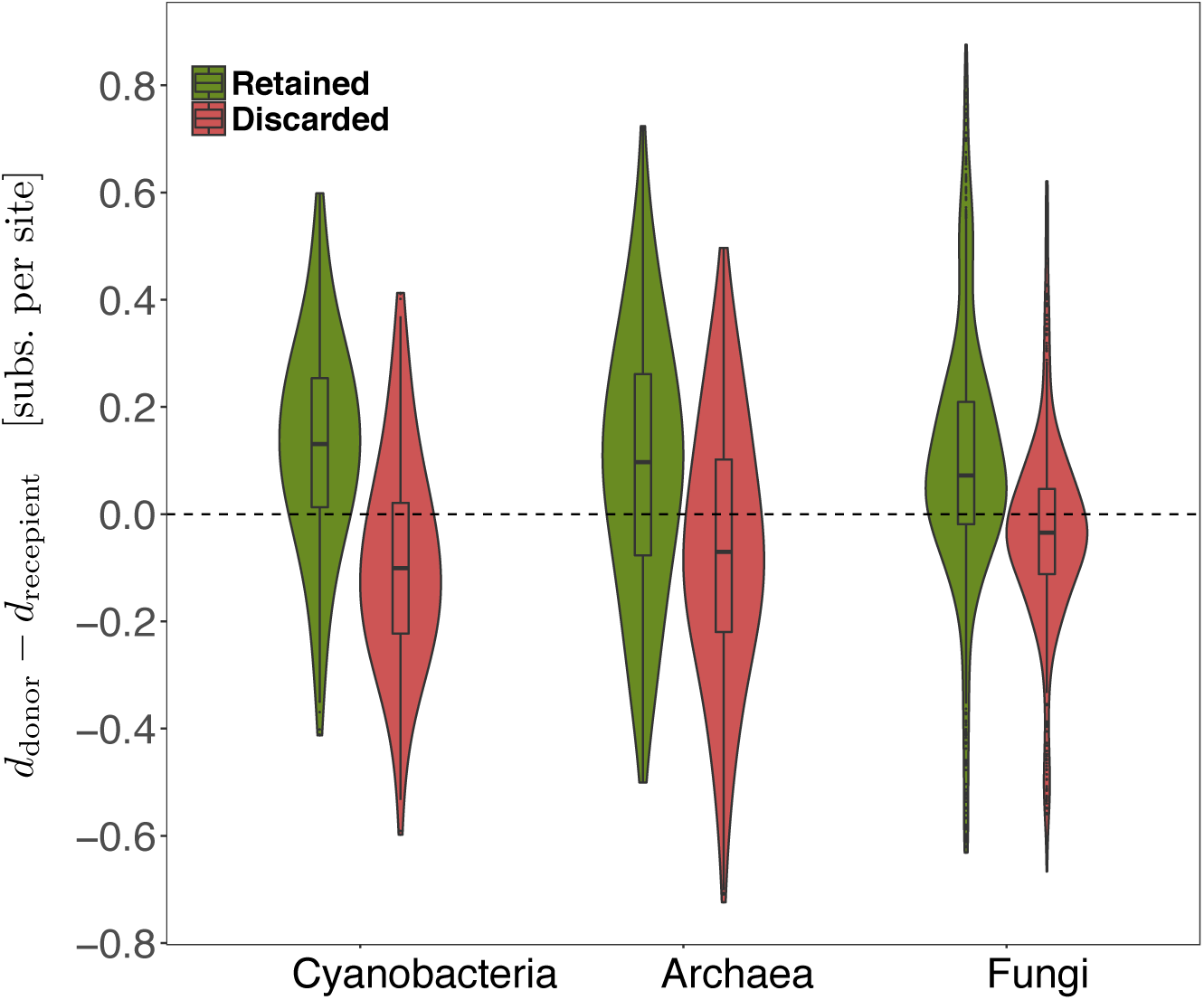
For genuine LGTs, the donor lineage must be at least as old as the recipient. As one proxy to investigate whether this was the case for transfers retained by our MaxTiC algorithm, we calculated clade-to-tip distances (see methods) for the inferred donor and recipient clades for LGTs that were retained and discarded by MaxTiC. (a) In all three datasets, transfers retained by MaxTiC (in red above) have the property that donor clades are further from the tips of the tree than recipient clades, but the opposite pattern is observed for conflicting transfers rejected by MaxTiC (green above), consistent with the idea that MaxTiC distinguishes genuine LGTs from phylogenetic artifacts.

One obvious difference between fossil- and transfer-based relative ages in Fig. 2 is that the level of agreement is patently lower for the latter. While in mammals approximately half of the chronograms proposed by the lognormal model agree with 100% of relative constraints, for other datasets no model reaches 80% agreement. This means that some relative constraints derived from LGT consistently disagree with uncalibrated molecular clock estimates. These disagreements are difficult to interpret because both molecular clocks and our transfer-based inferences may be subject to error; simulations suggest that spurious gene transfer inferences do occur with ALE, albeit at a low rate (Chauve et al.^35^, Fig. S23). Nonetheless this low error rate on simulations suggests that at least some transfers contradicting the molecular clocks are genuine. This yields the exciting idea of a new source of dating information, independent of and complementary to the molecular clock. To gain further insight into the robustness of these transfer-based estimates, we evaluated their statistical support from the data. Since MaxTiC yields a fully ordered species tree, the relative age constraints derived from its output are potentially overspecified and include constraints with relatively low statistical support. To ascertain the extent of overspecification, we evaluated the statistical support of relative constraints by taking random samples of 50% of gene families and reconstructing the corresponding MaxTiC 1000 times (Figs. S20-S22). We then counted the number of times a constraint was observed. In all datasets, a large majority of constraints were highly supported (found in at least 95% of the replicates) and among these, a significant number (between 20% and 32%) consistently disagreed with molecular clock estimates (see Table S2). These strongly-supported transfer-based constraints that disagree with the clocks could result from the inability of uncalibrated molecular clock estimates to recover the correct timing of speciations in groups with large variations in the substitution rate over time.

Specifically, LGTs provide strong support for the relatively recent emergence of the Prochlorococcus - Synechococcus clade in Cyanobacteria (blue clade in Fig. 4a), irrespective of uncertainty in the root of Cyanobacteria (see Supp. Mat.). Although the Prochlorococcus - Synechococcus clade is inferred to be ancient by three of the four uncalibrated molecular clock models in our study, previous analyses using relaxed molecular clock methods with more extensive species sampling and several fossil calibrations, including fossils dating akinete forming cyanobacteria at ∼2.1 Gya^36^ (green in Fig. 4.a) have consistently dated this clade as younger than most of the rest of cyanobacterial diversity^37,38^. Prochlorococcus have a known history of genome reduction and evolutionary rate acceleration^23^, which may lead to artifactually ancient age inferences under uncalibrated molecular clock models, as for rodents above. This demonstrates that relative time orders implied by LGT can, like fossils, provide a consistent dating signal that is independent of the rate of sequence evolution.

**Figure 4.**
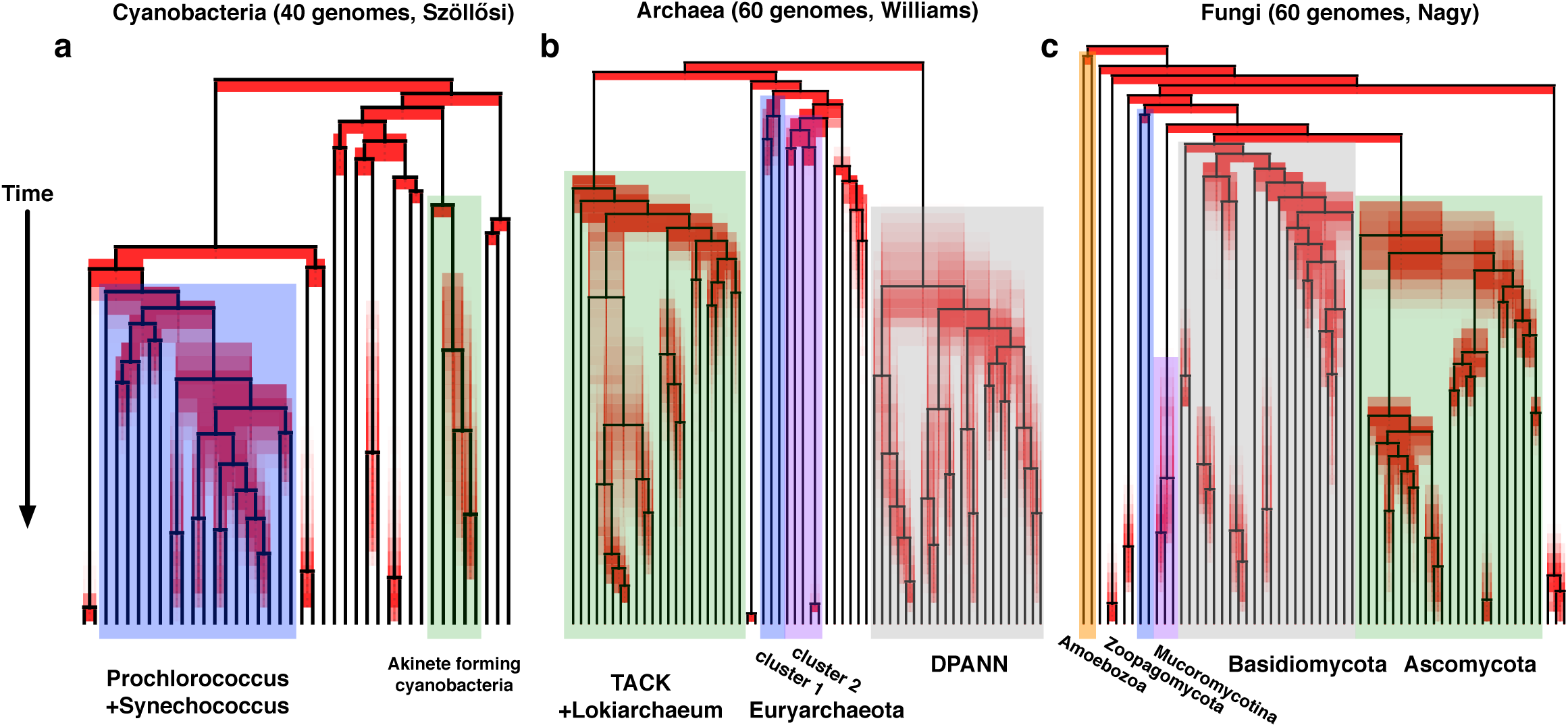
The order of speciations according to LGT. 5000 chronograms with a speciation time order compatible with LGT-based constraints were sampled per data set (a: Cyanobacteria, b: Archaea, c: Fungi). The black line corresponds to the consensus chronogram. Red shading represents the spread of node orders within the sample: nodes are in bright red if there is little or no uncertainty on their order according to LGT, in a light red smear if there is high uncertainty on their order. Clades discussed in the text are labelled and shaded. Supplementary Figures S1, S2 and S3 provide the same consensus chronograms with species names at the tips.

In Archaea, patterns of LGT suggest that several nodes within the Euryarchaeota including cluster 1 and 2 methanogens (blue and purple clades in Fig. 4) are older than both the TACK+*Lokiarchaeum* clade (red clade in Fig. 4, the clade uniting Thaumarchaeota, Crenarchaeota, Aigarchaeota, Korarchaeota and *Lokiarchaeum*) and the DPANN Archaea (grey in Fig. 4, a genomically diverse group with small cells and genomes, with reduced metabolism suggestive of symbiont or parasite lifestyles). The relative antiquity of methanogens is consistent with evidence of biogenic methane at a very early stage of the geological record (∼3.5 Gya^39^), and with another recent analysis that used a single LGT to place the origin of methanogens before the radiation of Cyanobacteria^14^. These relationships are not recovered by any of the molecular clock models, and suggest that LGT-derived constraints may be highly informative for future dating studies.

The relative order of appearance of archaeal energy metabolisms corresponds to increasing energy yield, with methanogenesis evolving before sulphate reduction, and the oxidative metabolisms of Thaumarchaeota and Haloarchaea evolving most recently. In addition, we find that *Ignicoccus hospitalis* branches before its obligate parasite *Nanoarchaeum* (cf. Fig. S2), despite the early divergence of the DPANN clade from other Archaea.

In Fungi, we recover LGTs that provide information on the order of some of the deepest splits. In particular, among crown groups, LGTs indicate that Zoopagomycota^40^ (blue in Fig. 4) diverged earlier than Mucoromycotina, Basidiomycota and Ascomycota (purple, grey and green in Fig. 4). Note that some inferred LGTs could result from processes such as hybridisation or allopolyploidisation, and that these processes contribute dating information that can be treated in the same way as LGTs. On a wider scale, between Eukaryotic groups, LGTs suggest that Amoebozoa (the outgroup, yellow in Fig. 4) diversified earlier than Opisthokonta and Apusozoa (the ingroup). This indicates that LGTs could strongly reduce the uncertainty associated with the divergence of the major eukaryotic clades^41^.

Our demonstration that clocks and transfers contain complementary and compatible dating signals casts the phylogenetic discord of LGT in a new light. This calls for the development of new methods to combine relative age constraints derived from transfers with molecular clocks and fossils calibrations. In principle, such a method can be developed by extending current Bayesian relaxed molecular clocks to include a prior on relative age constraints, as is now done with fossil calibration priors. An alternative approach would be to jointly infer a dated species tree together with thousands of dated and reconciled gene trees in the context of a hierarchical probabilistic model^47^; this approach would be theoretically superior, but likely computationally prohibitive at the present time^47^.

The geological record of microbial life is sparse, and its interpretation is fraught with difficulty. Our results show that there is abundant information in extant genomes on dating the Tree of Life waiting to be harvested from the reconstruction of genome evolution. This signal mostly contains information on the relative timing of diversification of groups that have exchanged genes through LGT, but we foresee several strategies to relate this relative timing to the broader history of life on Earth. First, gene transfers between bacteria and multicellular organisms that have left a trace in the fossil record will allow the propagation of absolute time calibrations to the microbial part of the Tree of Life^42^. Similarly, the signal of coevolution between hosts and their symbionts, such as in the gut microbiome of mammals^43^, could also be used to propagate absolute dating information from the host to the symbiont phylogeny. Finally, geochemistry can provide major constraints on early evolution^44,45^: for example, LGT events to the ancestors of bacteria capable of oxygenic photosynthesis, *i.e.* Oxyphotobacteria^46^, imply that the donor lineages must be older than the oxygenation of Earth’s atmosphere at approximately 2.3 Gya^44,45^. Phylogenetic models of genome evolution have the potential to turn the phylogenetic discord caused by gene transfer into an invaluable source of information on dating the tree of life.

## Acknowledgements

G.J.Sz. received funding from the European Research Council (ERC) under the European Union’s Horizon 2020 research and innovation programme under grant agreement No. 714774. This project was supported by the French Agence Nationale de la Recherche (ANR) through grant no. ANR-10-BINF-01–01 ‘Ancestrome’. Computations were made using the Curie supercomputer thanks to PRACE project 2013081661 and the computing facilities of the CC LBBE/PRABI. T.A.W. is supported by a Royal Society University Research Fellowship. We thank N. Lartillot, T. Warnow, M. Paris, I. Derényi and J. Miguel Blanca Postigo for useful discussions, comments on the manuscript and further computing resources.

